# Replication-Enhanced Detection of Quantitative Traits Evolving Adaptively (REDQuanTEA): an improved statistical framework to detect locally adaptive traits

**DOI:** 10.64898/2026.08.31.748449

**Authors:** Siyuan Feng, John E. Pool

## Abstract

Comparing quantitative trait differentiation (*Q*_ST_) with neutral genetic differentiation (*F*_ST_) is an established approach to detect locally adaptive trait differentiation, but empirical applications can lose power when *Q*_ST_ is deflated by extrinsic trait variance (i.e. from non-genetic sources such as environmental effects and measurement error). We present REDQuanTEA (Replication-Enhanced Detection of Quantitative Traits Evolving Adaptively), a ready-to-use computational workflow that leverages biologically replicated data to disentangle genetic and extrinsic trait variance, while using Approximate Bayesian Computation (ABC) to refine estimates of *Q*_ST_. Instead of comparing all traits with a single *F*_ST_-based cutoff, REDQuanTEA generates trait-specific dynamic outlier *Q*_ST_ thresholds from neutral *F*_ST_ distributions based on matching experimental properties and the trait-specific level of extrinsic variance. In addition to identification of candidate adaptive traits from empirical data, the package enables simulation-guided assessment of experimental design and statistical analysis options. Using a demographic benchmark based on *Drosophila melanogaster*, REDQuanTEA outperformed estimators based on analysis of variance (ANOVA), particularly when extrinsic variance was moderate to high, while controlling false positive rates (FPRs). Assuming fixed experimental effort, two replicates with more independent genotypes often outperformed three replicate designs when extrinsic variance was low, and performed similarly as extrinsic variance increased. REDQuanTEA therefore provides a framework for optimizing experimental plans and detecting adaptively differentiated traits with improved power.

## 1 Introduction

Trait differentiation between populations can be shaped by drift, migration, mutation, and selection. A classic way to test whether genetically based trait differentiation exceeds neutral expectations is to compare quantitative trait differentiation (*Q*_ST_) with putatively neutral genetic differentiation (*F*_ST_) (Wright, 1949; Weir and Cockerham, 1984; Lande, 1992; Spitze, 1993; Merilä and Crnokrak, 2001; McKay and Latta, 2002; Leinonen et al., 2013). Among neutrally evolving additive traits, mean *Q*_ST_ is expected to equal mean *F*_ST_ at neutral loci. Individual trait *Q*_ST_ values that fall above the distribution of neutral values are therefore interpreted as evidence for directional, locally adaptive differentiation, whereas *Q*_ST_ values within the neutral distribution are consistent with uniform or stabilizing selection across populations (Lande, 1992; Spitze, 1993; McKay and Latta, 2002; Leinonen et al., 2013).

Despite this clear conceptual framework, empirical *Q*_ST_–*F*_ST_ analyses are often limited by how trait variance is measured. Trait values encompass genetic effects as well as “extrinsic” (non-genetic) effects from rearing conditions, sample handling, and measurement error. If this extrinsic variance is not estimated separately, such environmental and experimental influences can inflate within- and between-population variance terms, which will tend to reduce the relative difference between them and therefore deflate *Q*_ST_ estimates on average, thus reducing power to detect adaptive trait differentiation (O‘Hara and Merilä, 2005; Whitlock, 2008; Huang et al., 2021). Common-garden and breeding designs can reduce non-genetic variance, but rigorous half-sibling or similar designs are laborious and can be infeasible for many study systems (Whitlock, 2008; de Villemereuil et al., 2022). Even in controlled experiments, measurement effects inevitably remain in the observed trait values.

A related problem is limited comparability of *Q*_ST_ across traits with different levels of extrinsic variance. This issue is especially important for molecular traits, which can differ markedly in extrinsic variance profiles. For example, transcript abundance is often noisier than downstream protein abundance (Liu et al., 2016). If two traits have the same estimated *Q*_ST_ but very different extrinsic variance, they do not provide the same strength of adaptive evidence, and so without explicit correction, cross-trait comparisons can be misleading. The third problem is that estimates of *F*_ST_ are unlikely to equal estimates of *Q*_ST_ even for neutral loci and traits, because the sources of bias are different in these two measures and are hard to control (Whitlock, 2008).

Here, we present REDQuanTEA (Replication-Enhanced Detection of Quantitative Traits Evolving Adaptively) to address these challenges. First, REDQuanTEA utilizes genetically identical biological replicates (e.g. distinct individuals from inbred strains or crosses between them) to estimate and isolate extrinsic variance directly. REDQuanTEA also uses regression Approximate Bayesian Computation (ABC) to estimate trait *Q*_ST_. Instead of applying one universal *F*_ST_ cutoff to all traits, REDQuanTEA generates trait-specific neutral *Q*_ST_ thresholds from neutral *F*_ST_ input, which improves power to detect adaptive traits, enhances cross-trait comparability, and calibrates comparisons between trait *Q*_ST_ and neutral expectations. In addition to these statistical advantages, REDQuanTEA is distributed as an automated and reproducible computational workflow. It can analyze tens of thousands of traits from a single command on either local Linux machines or high-throughput computing systems.

## 2 Materials and Methods

### 2.1 Overview of REDQuanTEA workflow

REDQuanTEA is designed to test whether quantitative traits show evidence of local adaptation while accounting for non-genetic environmental and measurement noise (extrinsic variance). The workflow requires three core inputs: (i) a sample structure table specifying the population, individual, and replicate membership of each sample; (ii) trait values measured on the same replicated biological samples; and (iii) neutral *F*_ST_ for the focal population comparison, provided either as a distribution of *F*_ST_ values at neutral loci or as inputs from which REDQuanTEA can generate that distribution. The main detection output is a trait-level table reporting the ABC-estimated 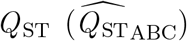, the trait-specific neutral threshold used for *Q*_ST_–*F*_ST_ comparison, and the adaptive call for each trait (Table S1). Optional benchmarking outputs summarize estimator performance across detection models (Table S2), levels of extrinsic variance (Table S2, Figure 3), and sample structures (Figure 4).

The major workflow of REDQuanTEA is shared across detection of adaptive traits and simulation-based evaluation of detection performance (Figure 1). First, a distribution of neutral *F*_ST_ values is obtained from direct input, VCF-derived allele counts, or coalescent simulation (Section 2.2). Second, empirical or simulated trait values are represented with the same nested population–individual–replicate sample structure (Section 2.3). Third, method-of-moments analysis of variance (ANOVA) decomposes trait variance into between-population genetic variance (*V*_*GB*_), within-population among-individual genetic variance (*V*_*GW*_), and extrinsic variance (*V*_*E*_) (Section 2.4). Fourth, regression ABC estimates 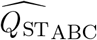 from observed summary statistics (Section 2.5). Finally, 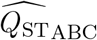 is compared with a dynamic neutral *Q*_ST_ threshold generated from neutral *F*_ST_ values and the relevant extrinsic-variance level (Section 2.6).

**Figure 1.**
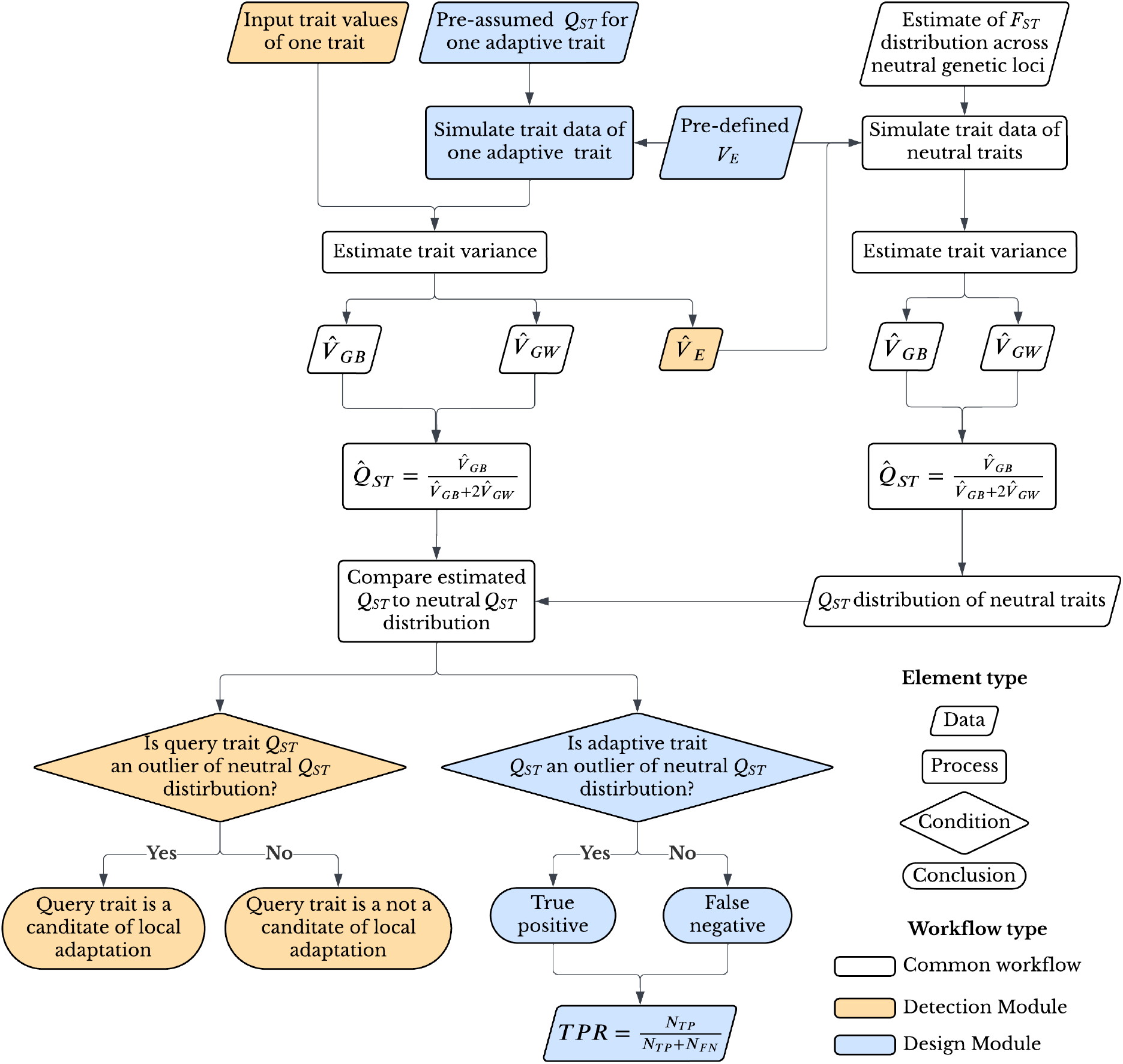
REDQuanTEA workflow for detecting adaptive traits and simulation-based benchmarking of detection performance. Neutral F_ST_ baselines are obtained from either user-supplied values, Variant Call Format (VCF)-derived allele counts, or coalescent simulations; replicated trait data are decomposed into genetic and extrinsic variance; ABC is deployed to estimate 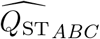; and dynamic neutral Q_ST_ thresholds identify candidate adaptive traits. A “Detection Module” detects adaptive differentiation of empirical traits, whereas a “Design Module” simulates trait data to evaluate model performance using the true positive rate (TPR).

These shared steps are implemented in two linked modules. The Detection Module applies the workflow to empirical trait data and detects traits that have undergone adaptive differentiation (Section 2.7). The Design Module simulates trait data across a spectrum of adaptive *Q*_ST_ values, processes them through the same estimation and thresholding steps, and quantifies detection performance (Section 2.8).

REDQuanTEA supports two types of running environments: it can run on a local Linux machine for small to medium batches of trait data, or it can run on HTCondor, a distributed computing system, for large trait panels or exhaustive summary statistic benchmarking.

### 2.2 Estimating the *F*_ST_ distribution of neutral genetic loci

REDQuanTEA uses a distribution of putatively neutral *F*_ST_ values as the genetic baseline for *Q*_ST_–*F*_ST_ comparison. This distribution can be supplied in three ways. In direct input mode, users provide one-column files of precomputed neutral *F*_ST_ values. In VCF mode, REDQuanTEA extracts allele counts for two user-specified populations from biallelic SNP genotypes, computes per-site *F*_ST_ values, and writes the same neutral *F*_ST_ files in direct input mode. In simulation mode, REDQuanTEA reads chromosome-specific ms parameters (Hudson, 2002), simulates neutral genotype matrices under the specified demographic model, computes per-site *F*_ST_ from simulated allele counts, and writes the same one-column neutral *F*_ST_ files. The distribution of neutral *Q*_ST_ values depends on the genetic architecture of the traits, with neutral variance maximized if traits are governed by a single locus. Hence, the use of single SNP *F*_ST_ values for the latter two options (and optionally for the first) corresponds to a conservative assumption of single locus trait architecture.

Both VCF-derived and simulation-derived paths use Reynolds’ estimator of the coancestry coefficient for genomic *F*_ST_ estimation (Reynolds et al., 1983). For site *ℓ*, REDQuanTEA calculates the numerator *a*_*ℓ*_ and denominator *a*_*ℓ*_ + *b*_*ℓ*_ as

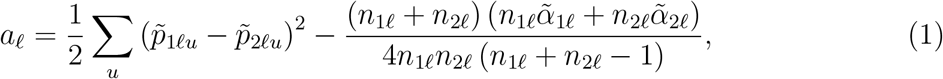

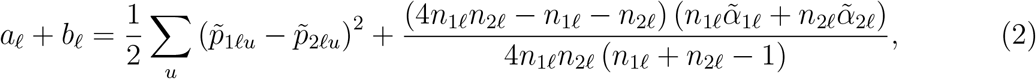

where 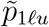 and 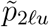 are the frequencies of allele *u* at site *ℓ* in the two populations, 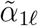 and 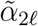 are the corresponding heterozygosities, and *n*_1*ℓ*_ and *n*_2*ℓ*_ are diploid sample sizes of each population at site *ℓ*. REDQuanTEA calculates per-site *F*_ST_ as

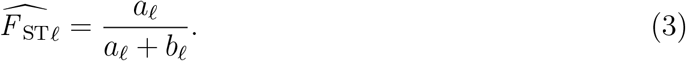

Under boundary cases where *a*_*ℓ*_ + *b*_*ℓ*_ = 0, 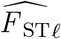 is set to zero. The current VCF parser skips multiallelic records, and users can also exclude such sites during preprocessing. Users should ideally restrict sites for *F*_ST_ estimation to neutral or nearly neutral genetic loci, such as four-fold synonymous sites in higher recombination genomic regions (if such rates of exchange are known), before interpreting the resulting *F*_ST_ distribution as a neutral expectation (although such sites may still be affected by selection at linked sites).

In simulation mode, the coalescent simulator ms (Hudson, 2002) is called with the user-provided mutation parameter, sample size, number of replicates, and demographic events such as population size changes, migration changes, and population mergers. One variable site is sampled from each non-recombining replicate. Each simulated segregating site is converted to allele counts in the two focal populations and passed through the same Reynolds estimator. The result is a neutral *F*_ST_ distribution that reflects the demographic scenario specified by the user. When autosomes and sex chromosomes differ in effective population size or demographic history, REDQuanTEA uses chromosome-specific neutral *F*_ST_ distributions for downstream *Q*_ST_–*F*_ST_ comparisons.

### 2.3 Sample structure and modeling of trait values

REDQuanTEA is built for replicated trait designs because replication makes extrinsic variance (*V*_*E*_) distinguishable from genetic variance, which is essential to avoid bias in *Q*_ST_ estimation. Such a design relies upon the possibility of measuring trait values from separate batches of genetically exchangeable individuals, such as from the same inbred strain, an outbred cross between inbred strains, or clonal individuals (hereafter “genotypes” or “individuals”). For empirical trait input, REDQuanTEA requires at least two populations, at least six genetically independent genotypes per population, and at least two biological replicates per genotype. In the basic two population setting, a trait value for population *p*, individual genotype *i*, and replicate *r* is modeled as

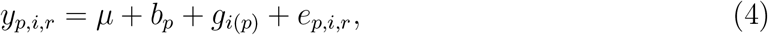

where *b*_*p*_ is the between-population genetic effect, *g*_i(*p*)_ is the within-population among-individual genetic effect, and *e*_*p,i,r*_ is the extrinsic effect, including environmental and measurement effects.

The same nested model is used to simulate trait data in REDQuanTEA’s Detection and Design modules. For two populations, REDQuanTEA parameterizes the between-population variance component *V*_*GB*_ by setting the population mean difference to

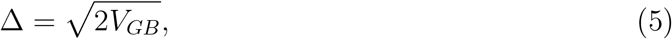

so that the two population means are *µ*_1_ = *µ* and *µ*_2_ = *µ* + Δ. Individual genotype effects and replicate-level extrinsic effects are then drawn as

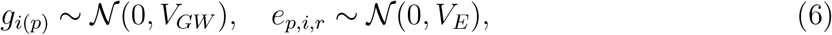

and simulated trait values are generated from the nested model above, with the same population, individual genotype, and replicate indexing as empirical input.

Importantly, REDQuanTEA compares focal *Q*_ST_ values not against neutral *F*_ST_ values, but instead against neutral *Q*_ST_ values simulated based on neutral *F*_ST_ values – thus accounting for error in *Q*_ST_ estimation. The variance parameters used for simulation are generated under two distinct contexts within the REDQuanTEA pipeline.

Under the first context, where REDQuanTEA simulates trait data sets from neutral *F*_ST_ values, it expects *Q*_ST_ = *F*_ST_ for each neutral locus. Because *Q*_ST_ is a function of between- and within-population additive genetic variance (Lande, 1992; Spitze, 1993),

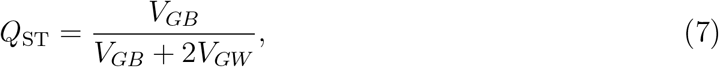

and total additive genetic variance in a common ancestral population can be written as (Leinonen et al., 2013, Box 1)

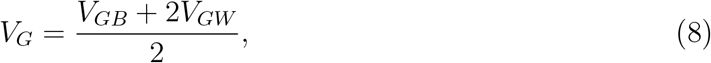

the genetic variance components can be derived as functions of neutral *F*_ST_ and *V*_*G*_:

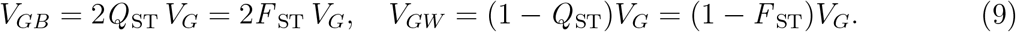

In the Detection Module, the accompanying extrinsic variance is derived from the ratio of the ABC-estimated *V*_*E*_ to the ABC-estimated *V*_*G*_ of empirical trait *t*:

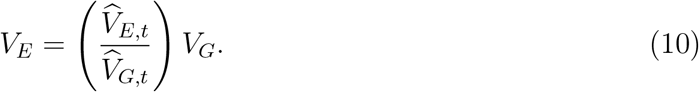

In the Design Module, *V*_*E*_ is set from predefined ratios of *V*_*E*_/*V*_*G*_ so that neutral and adaptive simulations can be evaluated under controlled noise conditions. Because *Q*_ST_ estimation depends on relative rather than absolute variance scales, REDQuanTEA sets *V*_*G*_ = 1 for simplicity when computing *V*_*GB*_, *V*_*GW*_, and *V*_*E*_ (Eqs. 9–10). Trait values can then be simulated under the nested model from the calculated variance components (Eqs. 4–6).

Under the second context of ABC estimation of variance components, variance parameters are drawn from their corresponding prior distributions and used to generate simulated data sets. These prior-drawn parameters are used to simulate trait data sets for ABC estimation of either an empirical trait in the Detection Module or a simulated trait in the Design Module. Specifically, for ABC simulation *i*, REDQuanTEA draws variance parameters as standard deviations *σ*_*GB*_, *σ*_*GW*_, and *σ*_*E*_:

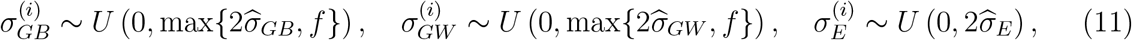

where 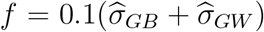 prevents priors of genetic variances from collapsing to identical or nearly identical values when ANOVA-estimated genetic variances are close to zero, which undermines ABC estimation.

### 2.4 Decomposition of trait variance

Using method-of-moments ANOVA estimators from replicated trait data, REDQuanTEA estimates extrinsic, within-population genetic, and between-population genetic variance components as

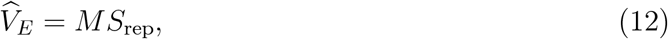

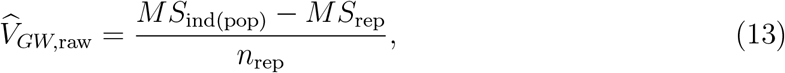

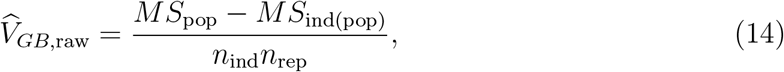

where *MS* terms indicate mean squared deviations from the mean, *n*_ind_ is individual genotypes per population and *n*_rep_ is replicates per genotype.

Method-of-moments estimates of 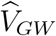 or 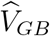 are unconstrained and can be negative or zero, especially if *V*_*E*_/*V*_*G*_ is high, but such estimates are unusable for calculating 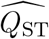. A classical fix would set each negative component to zero (Searle et al., 1992), but since 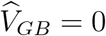 yields 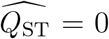 and 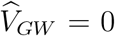 yields 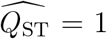, the resulting distribution could become artificially enriched for extreme values. So instead, REDQuanTEA adds a smooth numerical stabilizer to raw 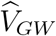 or 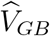. Let

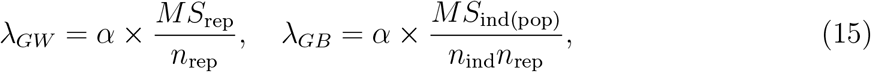

with default *α* = 0.1 (*λ* scaled to the ANOVA noise that enters each component). Each raw variance component is modified as

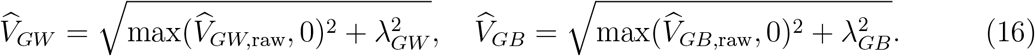

When extrinsic variance ratio is low, 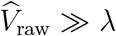 and 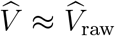; when extrinsic variance ratio is high and 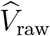 is close to 0, *Q*_ST_ will be pushed away from 0 or 1.

Although *Q*_ST_ can be calculated directly from the above modified ANOVA-estimated variance components, REDQuanTEA uses these ANOVA estimates primarily as summary statistics of trait data and estimates final *Q*_ST_ with ABC.

### 2.5 *Q*_ST_ estimation by regression ABC

ABC bases *Q*_ST_ inference on parameter values that could have generated features of the observed data, instead of relying on ANOVA point estimates. While replication is expected to allow ANOVA to separate extrinsic variance (*V*_*E*_), ANOVA estimates of *V*_*E*_ may fail to separate all extrinsic variance from ANOVA estimates of *V*_*GW*_ and *V*_*GB*_, especially when true *V*_*E*_ is high compared to genetic effects. In part because *Q*_ST_ depends on a ratio involving *V*_*GW*_ and *V*_*GB*_ (Eq. 7), direct ANOVA-based *Q*_ST_ can be noisy (O‘Hara and Merilä, 2005; Whitlock, 2008).

For each observed trait or simulated observed data set, REDQuanTEA’s ABC approach samples parameter values from prior distributions, simulates trait data using those prior parameters (Section 2.3), computes the same summary statistics for each simulated data set, and calculates a Euclidean distance from the observed summary statistics. After ranking simulations by distance, a tolerance rate (default: 0.001) is applied to retain the closest simulations for regression adjustment. During regression adjustment of retained parameter values, closer simulations receive higher weight according to their distance from the observed summaries. The adjusted parameters of retained simulations approximate the posterior parameter distributions for estimating trait *Q*_ST_.

REDQuanTEA uses local linear regression correction by default, following the regression-adjustment ABC approach of Beaumont et al. (2002) as implemented in the abc framework (Csilléry et al., 2012). For each retained, regression-adjusted posterior draw *k* = 1, …, *K*, REDQuanTEA calculates *Q*_ST_ using the adjusted variance components and reports their posterior mean:

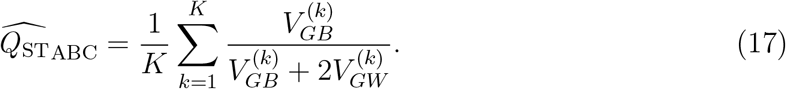

ABC summary statistics also include *V*_*GB*_, *V*_*GW*_, and *V*_*E*_, as well as derived ratios and partition fractions (Table 1). The performance of ABC models with different combinations of summary statistics is benchmarked in the Design Module (Section 2.8, Table S2).

**Table 1:** ABC summary statistics used by REDQuanTEA. Here V_GB_ and V_GW_ denote between-population and within-population genetic variance components, respectively, and V_G_ = V_GB_ + V_GW_. Only a subset of these statistics is typically used simultaneously.

| Summary statistic | Definition |
| --- | --- |
| $SD_{GB}$ | $\sqrt{V_{GB}}$, the standard deviation corresponding to between-population genetic variance. |
| $SD_{GW}$ | $\sqrt{V_{GW}}$, the standard deviation corresponding to within-population genetic variance. |
| $SD_E$ | $\sqrt{V_E}$, the standard deviation corresponding to extrinsic variance among replicates. |
| $Q_{ST}$ | Quantitative trait differentiation. |
| $V_E/V_G$ | Extrinsic variance relative to total genetic variance. |
| $V_{GB}/V_{GW}$ | Between-population genetic variance relative to within-population genetic variance. |
| $V_{GW}/V_E$ | Within-population genetic variance relative to extrinsic variance. |
| $V_{GB}/V_E$ | Between-population genetic variance relative to extrinsic variance. |
| $V_{GB}/(V_G + V_E)$ | Between-population genetic variance relative to total genetic plus extrinsic variance. |
| $V_{GW}/(V_G + V_E)$ | Within-population genetic variance relative to total genetic plus extrinsic variance. |
| $V_E/(V_G + V_E)$ | Extrinsic variance relative to total genetic plus extrinsic variance. |

### 2.6 Comparing *Q*_ST_ with a dynamic neutral *Q*_ST_ threshold

REDQuanTEA detects adaptively differentiated traits through *Q*_ST_–*F*_ST_ comparisons. In general, it is recognized that a trait’s *Q*_ST_ value should be compared against a full neutral *F*_ST_ distribution (or for the present method, the neutral *Q*_ST_ distribution derived from it), rather than comparing a given 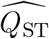 value against a single 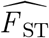 or the mean 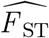 across loci. Just as neutral genetic loci yield a distribution of *F*_ST_ values due to random evolutionary variance, the loci underlying neutral traits will vary somewhat in their degree of differentiation, with this variance in neutral *Q*_ST_ values being maximized under a single locus genetic architecture as conservatively assumed here. In addition to comparing a focal *Q*_ST_ value against a neutral distribution (of *Q*_ST_ values), REDQuanTEA incorporates a dynamic outlier threshold based on the estimated level of extrinsic variance, along with estimator calibration. For a trait or simulation condition *t* with neutral *F*_ST_ values 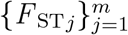 and relevant extrinsic variance 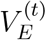 (Section 2.3), REDQuanTEA simulates neutral trait data sets under each *F*_ST*j*_, re-estimates 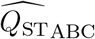 with the same ABC model used for the observed trait or adaptive simulation (Section 2.5), and obtains the calibrated neutral distribution

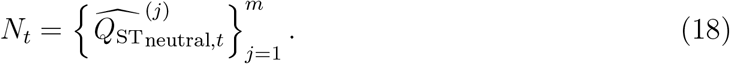

*T*_*t*_ is the threshold value corresponding to the user-specified quantile level *α* (default: *α* = 0.95) of the neutral distribution *N*_*t*_,

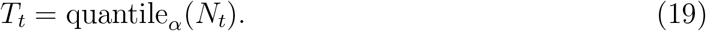

In the Detection Module, 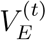 is estimated from the empirical trait by ANOVA. In the Design Module, 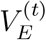 is supplied as a predefined extrinsic-variance level. In both modules, *T*_*t*_ is therefore dynamically determined by 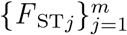 and 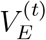.

Trait differentiation is labelled as potentially adaptive when the focal *Q*_ST_ estimate exceeds its calibrated threshold, e.g., 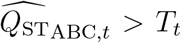 for empirical trait *t* in the Detection Module. Thus, the comparison is essentially between 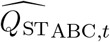 and *N*_*t*_ generated under matching sample structure, estimator, and extrinsic variance level. Rather than comparing all trait *Q*_ST_ values with one universal neutral *F*_ST_ distribution, dynamic *Q*_ST_-based thresholding compares each trait with a calibrated neutral baseline, which accounts for context-specific estimator bias, enforces consistency of adaptive calls across traits with different levels of extrinsic variance, and fixes the false positive rate (FPR) at 1 − *α* for controlled evaluation of model performance.

One challenge is that traits governed by loci on chromosomes of different effective population sizes, such as autosomes and the X chromosome, will have differing neutral distributions of *Q*_ST_, because the distinct effective population sizes of these chromosomes alter the strength of genetic drift and therefore neutral *F*_ST_ distributions. Often it is not known a priori which chromosomes may underlie a given trait’s variation. However, in the case of -omic measurements associated with specific genes, an important contribution of cis-regulatory genetic variation might often be suspected, and therefore comparisons of such trait values with neutral *Q*_ST_ distributions corresponding to the chromosome of origin could be motivated. REDQuanTEA can therefore generate neutral *F*_ST_ baselines, neutral *Q*_ST_ distributions, and adaptive thresholds separately for chromosomes with differing effective population sizes.

### 2.7 Detection Module: Detection of adaptive trait differentiation

The Detection Module applies the shared REDQuanTEA workflow to empirical trait measurements. For each trait *t*, REDQuanTEA first joins the trait-value table to the sample structure table and calculates observed ANOVA summary statistics, including 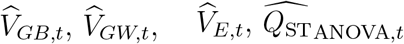, and other derived ratios (Table 1). These statistics summarize the observed trait data for ABC, and they also provide the trait-specific extrinsic variance estimate used for dynamic neutral threshold generation.

For empirical trait *t*, the final ABC estimate 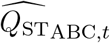 is obtained using the summary statistic combination selected by the user, who can also use results of the Design Module model bench-marking as a reference. For the same trait, REDQuanTEA then selects the chromosome-specific neutral *F*_ST_ distribution file based on the trait’s chromosome annotation. For each *F*_ST*j*_ in that file, the workflow simulates a neutral trait data set under the empirical sample structure, assumes that each neutral true *Q*_ST_ = *F*_ST*j*_, uses 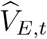, and re-estimates 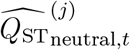 by the same ABC model. The resulting *N*_*t*_ for the empirical trait is compared with 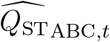 for adaptive calling (Section 2.6).

The final Detection Module output is a table with one row per trait. The table includes the trait identifier, chromosome type, 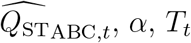, and a binary adaptive call that indicates whether 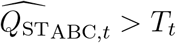 (Table S1). Because each trait receives its own *T*_*t*_ based on 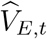, the Detection Module supports comparisons among adaptive calls of traits that differ in levels of environmental or measurement effects.

### 2.8 Design Module: Simulation-based evaluation of detection performance

The Design Module evaluates detection performance by simulating adaptive traits with known *Q*_ST_ and processing them through the same estimation and thresholding workflow used in the Detection Module. This module supports two purposes: benchmarking detection power of ABC models with different combinations of summary statistics to aid model selection, and benchmarking detection power of different sample structures to guide experimental design.

At the beginning of the Design Module workflow, REDQuanTEA simulates adaptive traits across a predefined spectrum of potentially detectable values *q*_*a*_. For each *q*_*a*_ in the adaptive grid (default full grid: *q*_*a*_ ∈ *{*0.50, 0.55, …, 1.00*}*) and for each extrinsic variance ratio *ρ* = *V*_*E*_/*V*_*G*_ ∈ *{*0.01, 0.1, 1, 10, 100*}*, the workflow derives *V*_*GB*_(*q*_*a*_) and *V*_*GW*_ (*q*_*a*_) from the *Q*_ST_ formula as described in Section 2.3, sets *V*_*E*_ from *ρ*, and simulates trait data sets under the nested population–individual–replicate model (Eqs. 4–6). Each simulated trait data set is then treated as if it were observed data: summary statistics are calculated, 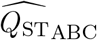 is estimated, and the estimate is compared with a neutral threshold.

Neutral thresholds in the Design Module are generated independently for each *ρ* and ABC model *c*. Let 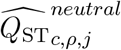 denote the 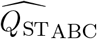 value for the neutral trait data set simulated from 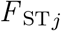. The neutral threshold is

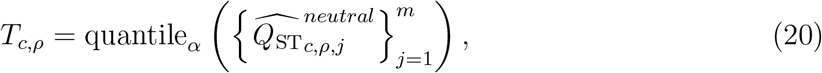

with *α* = 0.95 by default. For *R* replicates (default: *R* = 10,000) at true adaptive value *q*_*a*_, the TPR is calculated as

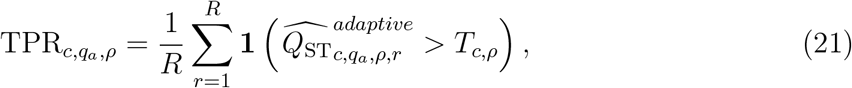

where 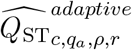 denotes the 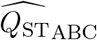 value for adaptive replicate *r*. The FPR is calculated from the neutral estimates as

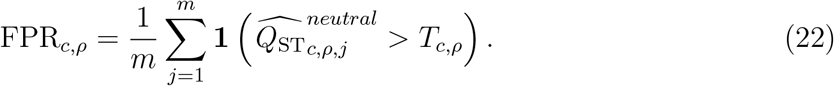

For a fixed sample structure and *ρ*, the model’s overall adaptive trait detection performance is summarized as the mean TPR across *q*_*a*_ values:

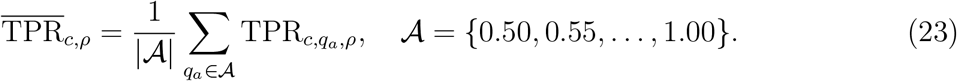

The outcome of the Design Module is a model-ranking table in which ABC models with different combinations of summary statistics are ranked by mean TPR (Table S2). As explained in Section 2.6, quantile-based thresholding fixes FPR at 1−*α*, allowing model performance to be summarized by TPR. The Design Module also supports sample structure comparison by repeating the same workflow on alternative numbers of genotypes and replicates. For a fixed *c* and expected *ρ*, this produces power comparisons across sample structures, as illustrated in Figure 4.

### 2.9 Benchmarking based on *Drosophila melanogaster* populations

For the empirical benchmarking in this manuscript, we used the neutral *F*_ST_ simulation strategy described in Section 2.2, with demographic parameters based on *D. melanogaster* populations (Sprengelmeyer et al., 2020). We used the coalescent simulator `ms` (Hudson, 2002) to simulate 100,000 independent neutral windows of 5,000 bp using chromosome-specific population mutation rates scaled by window length and demographic parameters describing population size changes, migration changes, and population mergers, as previously implemented (da Silva Ribeiro et al., 2022). We used Zambia and France populations as the focal comparison within the broader *D. melanogaster* demographic model and kept autosomal (based on arm 3L demographic estimates) and X-chromosomal simulations separate.

With the neutral *F*_ST_ distributions calculated from simulated *D. melanogaster* genotype matrices (Section 2.2), we used the Design Module to evaluate REDQuanTEA performance. First, we evaluated all 2,047 combinations of the 11 ABC summary statistics in Table 1 and ranked them by mean TPR, producing the model-ranking output reported as Table S2. We used this exhaustive end-to-end performance ranking for ABC model selection because there is no general analytic rule that identifies an optimal ABC summary-statistic model in advance. Sufficient summaries are usually unavailable for the complex models where ABC is most useful, and the best summary set can be data set- and target-specific (Joyce and Marjoram, 2008; Nunes and Balding, 2010; Fearnhead and Prangle, 2012; Blum et al., 2013).

Second, we assessed the TPR and FPR for detecting adaptive trait differentiation by comparing trait 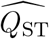 or 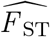 to dynamic thresholds based on neutral *Q*_ST_ distribution and fixed thresholds based on the raw neutral *F*_ST_ distribution (Figure 2). Third, we compared detection performance across extrinsic variance ratios and estimation methods, including for a top-ranked ABC model, ANOVA, and ANOVA on replicate-free data (Figure 3). The first, second and third analyses used a fixed sample structure including two populations, six genotypes per population, and three replicates per genotype, except that the second analysis also evaluated power for replicate-free data. Lastly, we ran sample structure comparisons for a top-ranked ABC summary statistic model to quantify how TPR changes with alternative allocations of genotypes and replicates (Figure 4).

**Figure 2.**
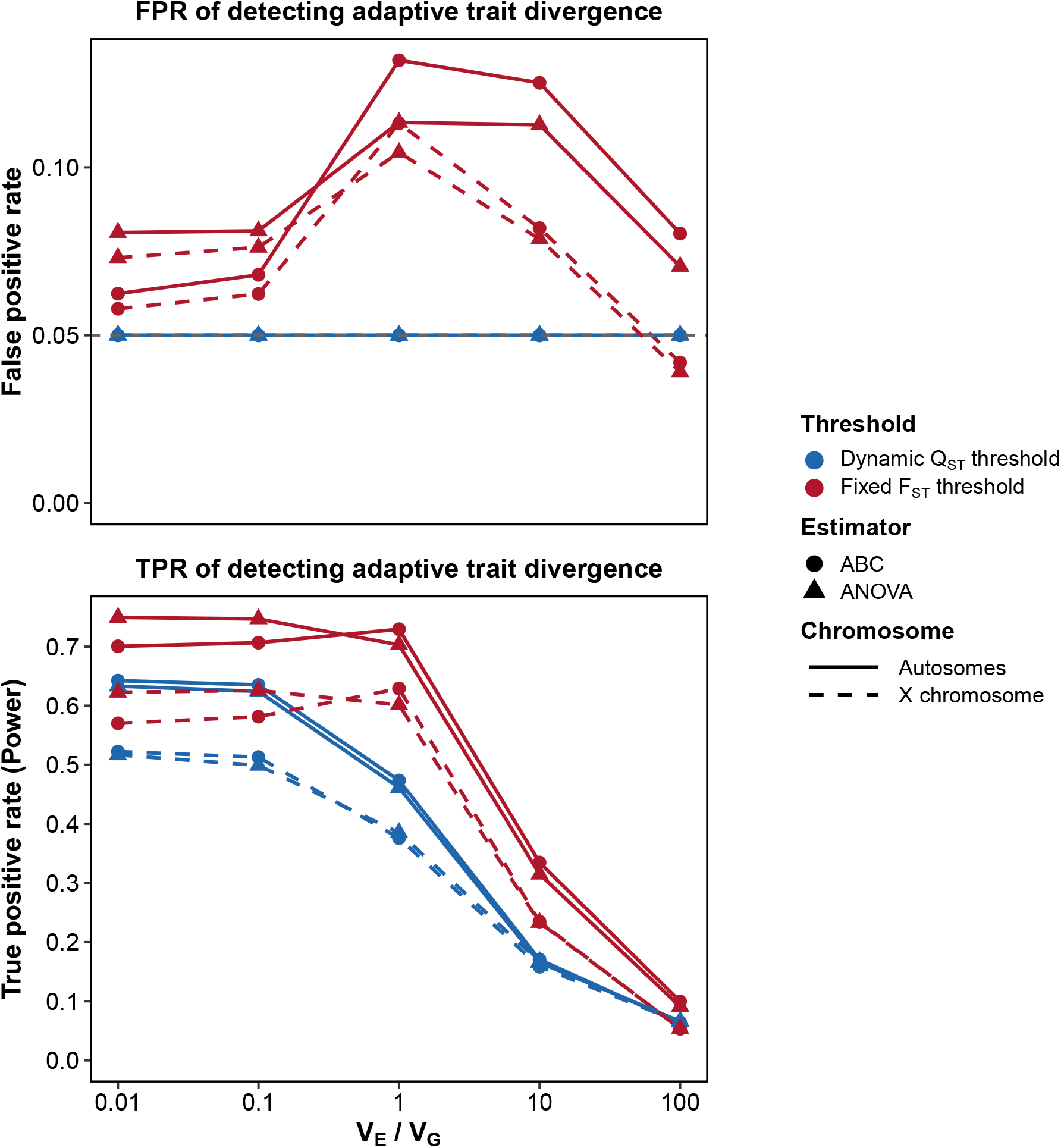
Unlike traditional comparisons using a neutral F_ST_ distribution, FPR to detect adaptive trait differentiation is controlled when using dynamic thresholding based on a neutral Q_ST_ distribution, with some cost to TPR. For a range of extrinsic variance ratios (V_E_/V_G_), results compare varying adaptive Q_ST_ re-estimated by ABC (dot) and ANOVA (triangle) to dynamic thresholds based on neutral Q_ST_ distribution (blue) and to fixed thresholds based on neutral F_ST_ distribution (red) using a D. melanogaster benchmark model, for autosomal (solid) and X-chromosomal simulations (dashed).

**Figure 3.**
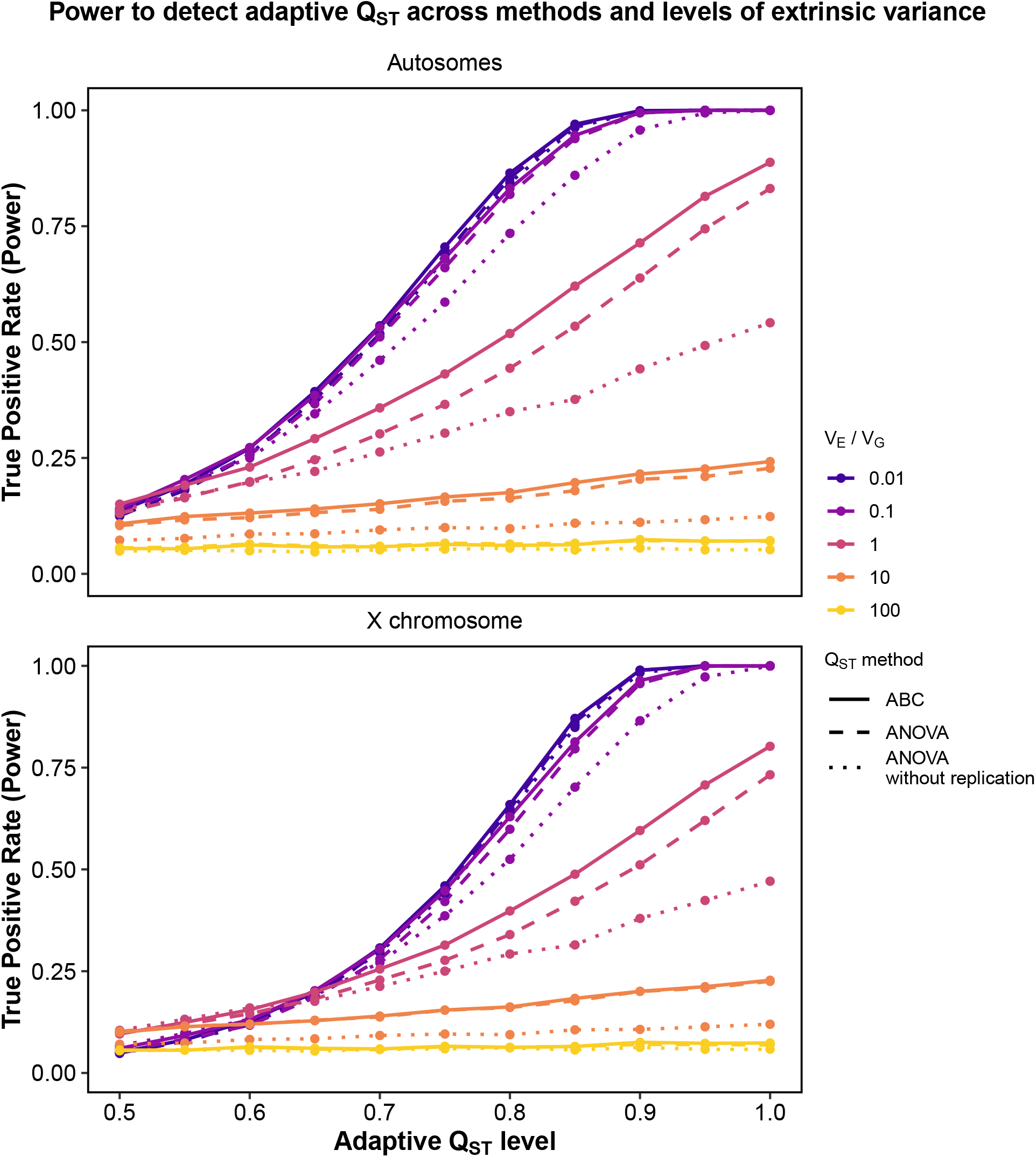
Power to detect adaptive differentiation is improved by replication, and in most cases by incorporating ABC estimation of Q_ST_. For a range of extrinsic variance ratios (V_E_/V_G_), results compare varying true adaptive Q_ST_ values to a neutral distribution based on the D. melanogaster benchmark model, for autosomal (above) and X-chromosomal simulations (below). The ABC Q_ST_ estimator uses replicated trait data and the default summary statistic combination; ANOVA is shown with and without replication for comparison.

**Figure 4.**
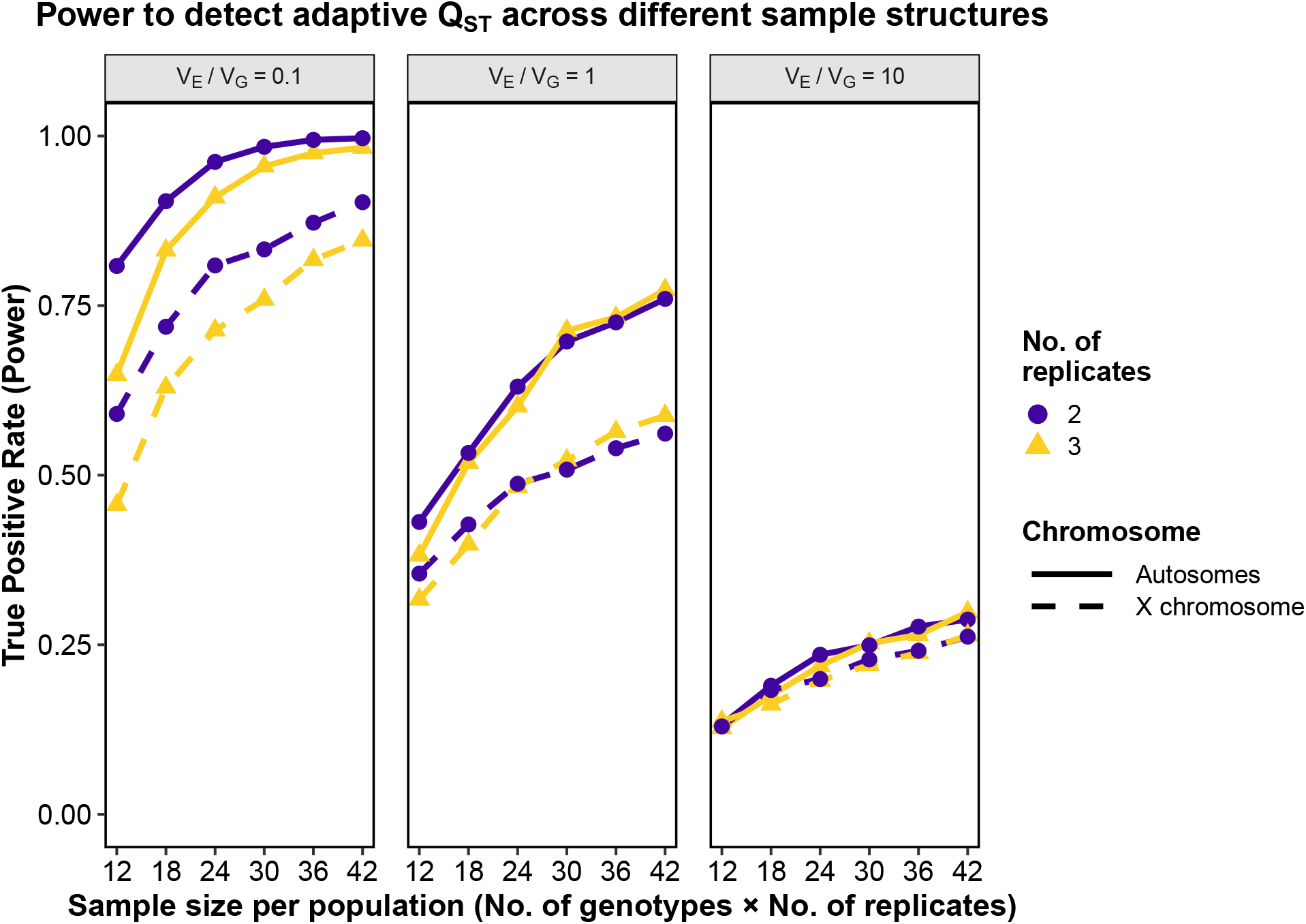
Experimental designs with two or three replicates have similar power to detect adaptive differentiation, except for an advantage of two replicate designs when extrinsic variance is relatively lower. Total experimental effort (number of samples) is the same between the two and three replicate designs in each case shown (the two replicate scenarios involve more unique genotypes instead). Results shown for three extrinsic variance ratios (V_E_/V_G_) are based on a true adaptive Q_ST_ of 0.82, a neutral distribution derived from the D. melanogaster benchmark (for autosomes and the X chromosome), and the REDQuanTEA default ABC summary statistic combination.

## 3 Results

### 3.1 Determinants of detection power for ABC summary statistic models

Using the Design Module and *D. melanogaster* models (Section 2.9), we benchmarked all 2,047 combinations of the 11 ABC summary statistics defined in Table 1; the full ranking is reported in Table S2. Mean TPR ranged from 0.2648 to 0.3645, but the top 100 combinations differed by only 0.0025, indicating that many high-ranking models performed nearly equivalently. Combination size was a major determinant of performance: larger summary statistic sets had lower mean TPR on average (Spearman *ρ* = −0.84), consistent with the known sensitivity of ABC to high-dimensional summary spaces (Blum, 2010; Blum et al., 2013).

Compact and weakly redundant models performed best. The best-ranked model contained one summary statistic, *V*_*GB*_/(*V*_*G*_ + *V*_*E*_) (mean TPR = 0.3645). The REDQuanTEA default, *Q*_ST_ and *V*_*GB*_/(*V*_*G*_ + *V*_*E*_), remained highly competitive (mean TPR = 0.3619); it keeps *Q*_ST_ as the biologically interpretable target while pairing it with a stabilizing partition fraction. Its difference from the best-ranked model was within stochastic simulation error, consistent with the negligible performance differences among other top-ranked ABC models.

### 3.2 Performance of REDQuanTEA to detect candidate adaptively differentiated traits

To benchmark REDQuanTEA’s capability to detect adaptive trait differentiation while avoiding false positives, we examined TPR and FPR while either using REDQuanTEA’s dynamic thresholding based on the neutral 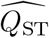 distribution or else using fixed thresholding based on the neutral 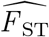 distribution (as commonly used in traditional *Q*_ST_–*F*_ST_ comparisons). Both 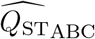 and 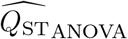 were compared against each type of the neutral threshold for adaptive calls (Figure 2). Compared to a FPR that was mechanistically stabilized at 1−*α* (*α* = 0.95 by default) using dynamic neutral 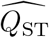-based thresholding, FPR were overall inflated when comparing trait 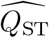 values, whether estimated by ABC or ANOVA, to a fixed neutral *F*_ST_-based threshold, because the 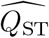-based threshold better controls for the error inherent in estimating *Q*_ST_ from a finite sample of individuals. Hence, REDQuanTEA’s conservative approach to avoid artificially high 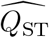 (Method section 2.4) controls FPR better than traditional *F*_ST_-based thresholds, at the cost of some power.

We then compared detection power across true adaptive *Q*_ST_ values, extrinsic variance ratios, chromosomes, *Q*_ST_ estimation methods, and replication structures (Figure 3). The ABC analyses used the REDQuanTEA default summary-statistic combination, *Q*_ST_ and *V*_*GB*_/(*V*_*G*_ + *V*_*E*_). As expected, adaptive traits with larger true *Q*_ST_ were easier to detect. At a moderate noise level where extrinsic variance equaled total genetic variance 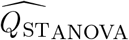 from replicate-free trait data had limited power. Adding replication immediately improved performance because replicated samples allow extrinsic variance to be estimated and separated from within-population genetic variance; otherwise, extrinsic variance inflates within-population variance and deflates *Q*_ST_. With the same replicated sample structure, switching from 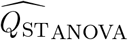 to 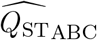 improved power again. This pattern motivates REDQuanTEA’s experimental and statistical framework: biological replication combined with ABC estimation, with an adaptive *Q*_ST_ cutoff defined by the trait-specific extrinsic variance estimate (dynamic thresholding).

The advantage of REDQuanTEA’s replicated design and ABC estimation depended on the relative amount of extrinsic variance. ABC provided an advantage over simple ANOVA when *V*_*E*_/*V*_*G*_ was 0.1, 1, or 10. The opposite trend was observed only when extrinsic variance was minimal (*V*_*E*_/*V*_*G*_ = 0.01). At the opposite extreme, no approach had meaningful detection power at *V*_*E*_/*V*_*G*_ = 100, due to minimal influence of genetic effects on trait variation. Thus, REDQuanTEA is most advantageous when extrinsic variance is large enough to bias standard *Q*_ST_ estimation but not so large that adaptive signal is almost completely obscured. These patterns were observed for both autosomes and the X chromosome.

### 3.3 Detection power varies by sample structure

To facilitate optimal experimental design, we compared how detection power changes under alternative allocations of individual genotypes and replicates, using a true adaptive *Q*_ST_ of 0.82 and the same default ABC model as in Section 3.2. These comparisons ask how to allocate a fixed sampling effort between sampling more independent genotypes versus adding more replicates per genotype.

Given a fixed total experimental effort, designs with two replicates and more genotypes performed better than those with three replicates with fewer genotypes when extrinsic variance was low (Figure 4). This pattern is expected because additional genotypes improve estimation of between-population and within-population genetic variance, whereas the third replicate mainly improves estimation of extrinsic variance (which in this case is low). When extrinsic variance was moderate or high, very similar results were observed between two and three replicate designs, indicating that benefits of either greater replication or additional genotypes were comparable.

These results suggest that replication should not be chosen in isolation. The best sample structure should depend jointly on the expected level of extrinsic variance, the neutral *F*_ST_ of the population pair, and the total sample budget. REDQuanTEA can therefore be used before data collection as a design tool: users can combine an expected neutral *F*_ST_ distribution, potential trait noise levels, and sampling budget to compare candidate allocations of genotypes and replicates under plausible *V*_*E*_/*V*_*G*_ values.

## 4 Discussion

### 4.1 Advances beyond standard *Q*_ST_–*F*_ST_ comparisons

REDQuanTEA makes the detection of adaptive trait evolution through *Q*_ST_–*F*_ST_ inference more feasible and powerful in a few ways. Existing *Q*_ST_–*F*_ST_ analysis is theoretically well motivated, but empirical use is often limited by extrinsic variance: uncontrolled environmental and measurement variance deflates *Q*_ST_ and reduces detection power, whereas rigorous breeding experiments can control this problem at a cost that could be infeasible for many organisms (O‘Hara and Merilä, 2005; Whitlock, 2008; Leinonen et al., 2013). REDQuanTEA addresses this limitation first through experimental design. By adding genetically identical biological replication to each studied genotype, it allows extrinsic variance to be estimated and separated from genetic variance, enabling accurate *Q*_ST_ estimation in broader study systems. REDQuanTEA then adds two statistical advances: ABC-based *Q*_ST_ estimation, which improves power over commonly used ANOVA-based estimators especially when traits have moderate or high extrinsic variance, and trait-specific outlier thresholds based on the extrinsic variance estimate, which makes *Q*_ST_–*F*_ST_ comparisons less biased and more comparable across traits with different levels of environmental and experimental noise. Furthermore, the practice implemented here of comparing empirical *Q*_ST_ values against an estimated neutral *Q*_ST_ distribution, rather than raw neutral *F*_ST_ values, can control false positives driven by uncertainty in the estimation of *Q*_ST_.

The REDQuanTEA workflow offers a framework that helps users before and after data collection. Before trait data collection, the Design Module can compare power estimates of candidate sample structures and guide the optimization of experimental design. The workflow also provides multiple ways to obtain neutral *F*_ST_ distributions, including direct input, VCF-derived estimates, and demographic simulation. After data collection, REDQuanTEA produces trait-level adaptive calls, benchmarking outputs, and reproducible records of the analysis. It uses an automated workflow implementation to minimize user effort and accommodates high-throughput trait input without compromising statistical performance.

More broadly, this *Q*_ST_–*F*_ST_-oriented framework allows us to ask a more evolutionarily refined question: not simply whether populations differ significantly, but whether neutral drift and population structure are sufficient to explain the observed trait differentiation. This distinction is especially important for molecular traits, such as from multi-omic data, where many among-population expression differences can reflect neutral drift or stabilizing selection as well as directional local adaptation (Whitehead and Crawford, 2006; Leinonen et al., 2013).

### 4.2 Considerations on experimental design

Beyond the individuals versus replicates allocation results described above (Section 3.3), the genetic material represented by each individual genotype can also affect *Q*_ST_ estimation and detection power. Inbred strains can be problematic for studies of trait variation, because by exposing rare recessive variation, inbreeding can inflate within-population variance (deflating *Q*_ST_) and make trait value distributions less representative of outbred population variation (Goudet and Büchi, 2006; Goudet and Martin, 2007; de Villemereuil et al., 2022). If sufficient inbred strains are available, one alternative is for each individual genotype to reflect the outbred F1 offspring of a cross between independent inbred strains.

### 4.3 Neutral *F*_ST_ distributions and genetic architecture of traits

The strongest neutral baseline is a demographic simulation or curated neutral locus set that matches the focal populations, chromosome class, sample size, and *F*_ST_ estimator. When the necessary demographic parameter estimates are unavailable, users can run REDQuanTEA’s empirical *F*_ST_ estimation with individual-level whole genome sequencing (WGS), restriction site associated DNA sequencing (RAD-seq; Baird et al., 2008), or genotyping by sequencing (GBS; Elshire et al., 2011) SNP data, and should subsample genotypes to match the diploid sample sizes of the phenotyped sample structure before estimating *F*_ST_ (such that *F*_ST_ and *Q*_ST_ sample sizes are the same). Chromosomes with different population histories or effective population sizes should be kept separate for neutral threshold generation.

Empirical genome-wide SNP *F*_ST_ distributions are usable but are not guaranteed to be neutral. If adaptively differentiated loci or linked selected regions (including linkage via inversions or low recombination regions) remain in the SNP set, the estimated “neutral” *F*_ST_ distribution can be inflated, raising the outlier threshold and reducing detection power. This bias is usually conservative for detecting *Q*_ST_ *> F*_ST_ outliers, but users should remove known selected loci, low recombination regions, and strongly linked marker clusters when such annotations are available (Edelaar et al., 2011; Cutter and Payseur, 2013; Li et al., 2019; da Silva Ribeiro et al., 2022).

An important conservative assumption inherent in this study’s use of neutral *F*_ST_ distributions from individual simulated or empirical SNPs is that the studied traits have a genetic architecture shaped by a single non-recombining locus. To the extent that traits have a multi-locus architecture, the expected neutral variance of *Q*_ST_ is reduced (due to averaging across multiple realizations of the evolutionary process). However, the genetic architecture of traits examined in evolutionary frameworks is generally unknown. Instead using *F*_ST_ distributions that assume a multilocus trait basis, or are drawn from longer recombining windows, should increase power but runs the risk of making an incorrect anticonservative assumption if some traits actually have a simpler genetic basis, and hence a broader neutral *Q*_ST_ distribution than assumed.

REDQuanTEA also inherits the central *Q*_ST_ assumption that genetic effects are additive (Lande, 1992; Spitze, 1993; Whitlock, 2008). In the replicated design, non-additive genetic effects do not automatically become extrinsic variance: dominance or epistatic effects that consistently differentiate individuals or populations can be absorbed into the among-individual or between-population genetic components, whereas environment-specific expression of those effects, including genotype-by-environment interactions among replicates, can inflate the extrinsic component. Because the neutral *Q*_ST_–*F*_ST_ expectation is derived for additive components, non-additive architectures can shift *Q*_ST_ values. Dominance often deflates *Q*_ST_ relative to *F*_ST_ under common island model assumptions. For epistasis, the simplest case of additive-by-additive epistasis is also expected to make neutral *Q*_ST_ lower than neutral *F*_ST_ on average. In both cases, tests for *Q*_ST_ *> F*_ST_ would be conservative (López-Fanjul et al., 2003; Goudet and Büchi, 2006; Goudet and Martin, 2007; Whitlock, 2008; Whitlock and Guillaume, 2009).

### 4.4 Future directions

In line with the above discussion on the importance of trait genetic architecture, a method able to integrate estimation of the number of contributing loci could boost power by allowing appropriate but narrower neutral *F*_ST_ and *Q*_ST_ distributions to be used. Such a method might need to incorporate additional sources of data, such as from other types of crosses (Castle, 1921).

There is a growing opportunity to apply evolutionarily informed analysis frameworks such as this one to the growing realm of -omic data types that quantify molecular traits. As replicated measurements of chromatin, transcript, protein, metabolite, and other molecular traits accumulate, REDQuanTEA can be optimized with trait-type-specific empirical expectations. For comparison between types of traits, one useful extension would be a trait-derived TPR correction for comparing the proportion of adaptive *Q*_ST_ outliers across trait types. For each trait type, REDQuanTEA could use the empirically estimated distribution of *V*_*E*_/*V*_*G*_ across observed traits to simulate neutral traits, generate matching trait-specific neutral *Q*_ST_ distributions, estimate TPR across adaptive *Q*_ST_ levels, and divide the observed outlier proportion by this trait-derived mean TPR. This extension would make outlier proportions more comparable across trait types that differ in extrinsic variance, such as mRNA expression and protein abundance. The same trait-derived simulations could also support an advanced model selection option that ranks ABC summary statistic combinations for a specific trait type, complementing the current general ranking strategy without assuming a fixed *V*_*E*_/*V*_*G*_ distribution.

Another direction is to extend *Q*_ST_–*F*_ST_ comparisons beyond pairwise population comparisons. While traditional *Q*_ST_–*F*_ST_ comparison is not limited to pairwise population comparisons (Spitze, 1993), and including more populations improves accuracy of between-population variance estimation and thus power of detecting spatially divergent selection (Whitlock and Guillaume, 2009), there are two major limitations with multi-population scenarios: the assumption of equal relatedness among the subpopulations that rarely holds in real-world can lead to false positives; the excess divergence cannot be localized to specific population contrast. The first limitation has been overcome by incorporating between- and within-population relatedness to model population structure (do Ó et al., 2025a,b). To locate adaptive trait divergence among multiple-populations, one route is to develop a quantitative trait analogue of branch-based genomic statistics such as the Population Branch Statistic (PBS; Yi et al., 2010) and Population Branch Excess (PBE; Yassin et al., 2016), in which trait differentiation on a focal population branch is compared with a neutral branch length expectation from two reference populations. A complementary route is a multi-population test that asks whether differentiation among trait values from multiple populations is compatible with the neutral coancestry matrix, rather than focusing on one comparison or population branch at a time. By continuing to develop methods such as these, the range of questions that can be asked about population trait differences in an evolutionarily informed framework can be expanded.

## Author Contributions

S.F. and J.E.P. designed REDQuanTEA and interpreted the results. S.F. implemented and benchmarked REDQuanTEA and wrote the manuscript. J.E.P. revised the manuscript.

## Acknowledgements

We thank members of the Pool laboratory for discussions on method design, benchmarking strategy, and workflow implementation. We especially thank Christopher McAllester, Michael Liou, and Qilin Li for providing consultation on statistical modeling.

## Data Availability Statement

REDQuanTEA source code, workflow documentation, and example input/output files are available at https://github.com/Sfeng666/REDQuanTEA. Benchmark settings used in this manuscript correspond to the public configuration files and commands in that repository.

